# Blocking cell fusion inhibits age-induced polyploidy and maintains epithelial organization in *Drosophila*

**DOI:** 10.1101/2021.08.09.455651

**Authors:** Ari S. Dehn, Navdeep Gogna, Patsy M. Nishina, Vicki P. Losick

## Abstract

A characteristic of normal aging and age-related diseases is the remodeling of a tissue’s cellular organization through polyploid cell growth. Polyploidy arises from an increase in nuclear ploidy or the number of nuclei per cell. However, it is not known whether age-induced polyploidy is an adaption to stressors or a precursor to degeneration. Here, we find that the adult fruit fly’s abdominal epithelium becomes polyploid with age through generation of large multinucleated cells that make up more than 40% of the tissue area. The syncytia arise by cell fusion, not endomitosis. Epithelial multinucleation is also a characteristic of macular degeneration, including *Ctnna1^tvrm5^,* a mouse model for pattern dystrophy. Similarly, we find that the knockdown of alpha-catenin enhances multinucleation in the fly epithelium. We further show that age-induced polyploidy can be suppressed by inhibiting cell fusion revealing a means to maintain tissue organization in older animals.

## Introduction

Aging is often accompanied by a permanent change to an organ or tissue’s cellular architecture, as aging cells turnover or undergo senescence. The lack of resident stem cell populations also limits the regenerative capacity of many animal tissues. As a result, cell growth via polyploidy may be the only mechanism to compensate for cell loss (Gjelsvik et al., 2019; Lazzeri et al., 2019). Polyploidy describes a cell that has more than the diploid copy of its chromosomes, hence referred to as 3C or greater chromatin content. Polyploidy occurs in a wide variety of cell types and organisms across the kingdoms of life, allowing cells to grow orders of magnitude larger as cell size scales with DNA content (Frawley and Orr-Weaver, 2015).

Studies in vertebrate and invertebrate models have demonstrated that polyploidy is essential for cellular adaption. These large cells speed wound closure and restore tissue mass when cell division is limited, protect against genotoxic stress, and promote resilience to environmental insults (Cao et al., 2017; Cohen et al., 2018; Grendler et al., 2019; Hassel et al., 2014; Losick et al., 2013; Storchova, 2014). Polyploidy is also associated with many disease states, in particular cancer. Tetraploidy predisposes cells to aneuploidy, whereas polyploidy in liver hepatocytes protects against tumorigenesis caused by the loss of heterozygosity (Fujiwara et al., 2005; Zhang et al., 2018). In the context of age and age-associated disease the biological significance of polyploidy is only beginning to be examined. In the mouse liver, hepatocytes are polyploid and further increase their ploidy with age (Donne et al., 2020). These polyploid hepatocytes have been found to repeatedly divide to maintain normal turnover of liver with age (Matsumoto et al., 2021). However, the accumulation of polyploid cells can affect liver metabolism, particularly when hepatocyte mitotic division is limited (Dewhurst et al., 2020). Likewise, neurons in both *Drosophila* and mammalian brain have also been found to increase ploidy with age. Tetraploid neurons are associated with Alzheimer’s disease and in the aging fly brain, polyploid neurons arise via the endocycle to protect against DNA damage (Lopez-Sanchez et al., 2017; Nandakumar et al., 2020).

Age-induced polyploidy has also been observed in mammalian eye tissues, including the cornea endothelium and retina pigment epithelium (RPE). Cornea endothelial cells are tetraploid and further increase ploidy in the age-associated disease Fuchs endothelial corneal dystrophy (Losick et al., 2016). RPE cells, which form an epithelial monolayer situated between the choroid blood supply and retina to form the blood retina barrier, has been observed to become multinucleated, hence polyploid, with age and in association with macular diseases, such as age-related macular degeneration (Chen et al., 2016; Saksens et al., 2016; Zhang et al., 2019). Additionally, RPE multinucleation has been reported in *Tvrm5*, a mouse model for Butterfly-shaped pigment pattern dystrophy as a pathological consequence of the disruption of *Ctnna1* (Saksens et al., 2016). Still the mechanisms controlling polyploidy in these cases, in particular with aging, remain unknown and importantly, whether these changes in tissue organization can be inhibited.

Here, we find that the adult *Drosophila* epithelium becomes polyploid with age by cell fusion, not failed cytokinesis. Blocking cell fusion via expression of a dominant negative Rac GTPase maintains the epithelial architecture of a young fly, which is primarily populated with mononucleated, diploid cells. The old fly multinucleated epithelium resembles the age-associated changes observed in mouse RPE. Indeed, we find that knockdown of conserved alpha catenin gene which is disrupted in Butterfly-shaped pigment pattern dystrophy also results in enhanced multinucleation in fly epithelium. Thus, our *Drosophila* model of age-induced polyploidy will be a useful adjunct to elucidate the genetic strategies utilized to maintain epithelial cell architecture with age.

## Results

### Polyploid cells arise by 20 days old in *Drosophila*

A previous study reported cell enlargement with loss of cell-cell junctions in the adult *Drosophila* ventral abdominal epithelium with age (Scherfer et al., 2013), but did not characterize the extent of multinucleation nor whether cells were becoming polyploid. Thus, we first characterized the extent of polyploidy by measuring epithelial cell size and number of nuclei per cell over a time course in days (d) from 5d to 50d post eclosion (Figure 1A-G). We used mated epi-Gal4 female flies for this study, which have a standard mean lifespan of 57d ± 4d (Figure 1H) (Koliada et al., 2020). In young flies (5d to 10d old), the epithelium was composed on average of 90% mononucleated cells with 5.6% binucleated and 2.28% multinucleated cells (Figure 1B, 1C, and 1I). By 20d old, there was a noticeable change in epithelial organization with large regions devoid of epithelial cell junctions and containing multiple epithelial nuclei (Figure 1D). There was also a significant increase in polyploid cells with more than doubling of binucleated and multinucleated cells. Polyploidy peaked in 40d old flies with 18% binucleated and 15% multinucleated epithelial cells (Figure 1F and 1I). The multinucleated cells in young flies contained 3-5 nuclei, whereas old flies had double the number of small multinucleated cells as well as frequent large multinucleated cells containing 6-31+ nuclei (Figure 1E-1G and 1J).

**Figure 1:**
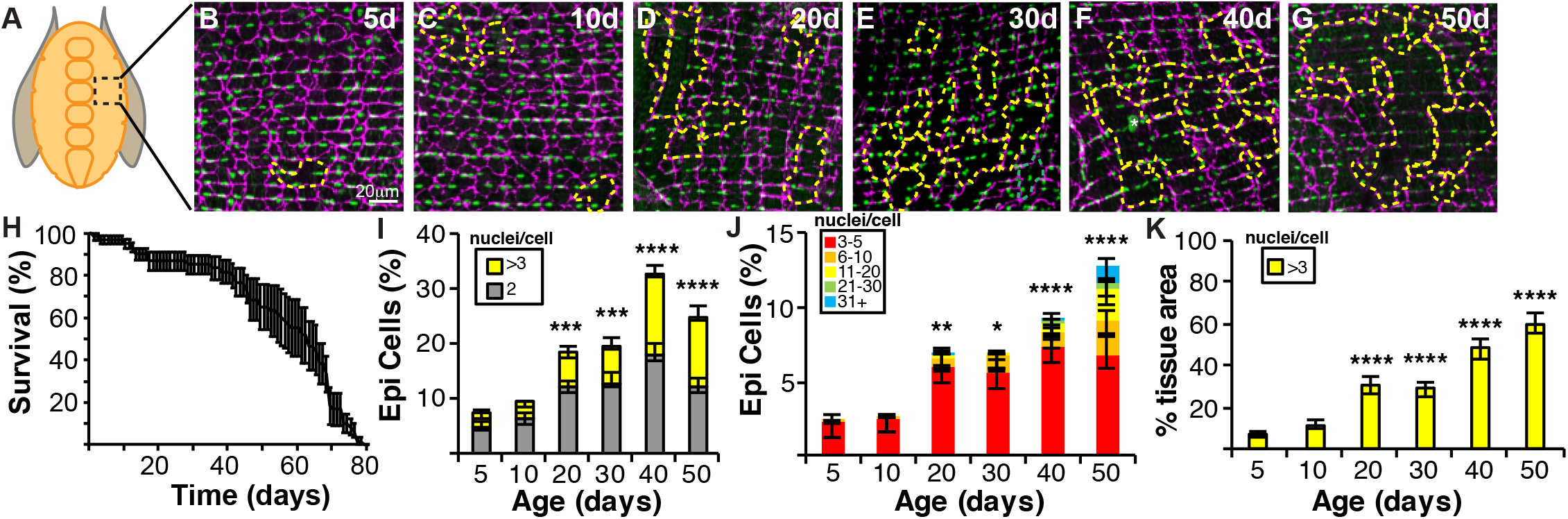
Age-induced polyploidy arises by 20 days in adult fruit fly epithelium. (A) Illustration of female *Drosophila* abdomen depicting the tissue region assayed. (B-G) Representative immunofluorescent images of the abdominal epithelium from epi-Gal4 flies aged 5d to 50d post eclosion. Septate junctions (FasIII, magenta), epithelial nuclei (Grh, green), and multinucleated cells (outlined, yellow dashed line). (H) *Drosophila* mated female survival curve. (I)Percentage of bi- and multinucleated epithelial cells with age. (J) Percentage of multinucleated epithelial cells by number of nuclei per cell. (K) Epithelial tissue area composed of multinucleated cells. Also, see Figure S1.

We noticed that the multinucleated cells (also known as syncytia), although few in number, made up the majority of tissue area with age. We calculated the percent of the epithelial area composed of syncytia and found the largest increase at 40d and 50d with multinucleated cells making up 48% and 60% of the tissue area, respectively (Figure 1K). Whereas, only 11% of the epithelial tissue area was made of multinucleated cells in 10d old flies. In conclusion, aging alters epithelial organization by generating multinucleated cells which affect the majority of the tissue area.

Next, we assessed epithelial organization in other *Drosophila melanogaster* strains to determine if the generation of syncytia with age was fly strain dependent. In young female *w^1118^* flies, we found that the epithelium resembled that of the epi-Gal4 strain and was composed of 89% mononucleated cells (Figure S1A and S1E). Similarly, in the old *w^1118^* strain, there was a significant increase in polyploidy and their epithelium was made up of 15% binucleated and 15% multinucleated cells (Figure S1B and S1E). Another wild-type *Drosophila* strain, Canton S, did have an elevated number of binucleated epithelial cells in young flies, but not multinucleated epithelial cells (Figure S1C and S1E). Yet by 40d old the Canton S strain showed a significant increase in its multinucleated cell population (Figure S1D and S1E). Thus, similar to epi-Gal4 fly strain, both the 40d old *w^1118^* and Canton S strains had at least 40% of the epithelium made up of multinucleated cells confirming that multinucleation is an aging-dependent, strain-independent phenomenon in *Drosophila melanogaster* (Figure S1F).

### Age-induced polyploidy is not dependent on apoptosis

As animals age, cells may not regenerate, thus leading to tissue deterioration. In *Drosophila,* we expected that syncytia to arise with age, similar to a wound healing response, to compensate for cell loss. To test this hypothesis, we measured changes in the cell and nuclear number in the epithelium from the aging time course (Figure 1A-G and Figure 2A-D). As expected, the total number of epithelial cells declined with age with a significant reduction at 20d old when there was corresponding increase in syncytia formed (Figure 2C). In total, 138 ± 5 cells made up a 22,500μm^2^ epithelial region which was reduced to 50 ± 5 cells in 50d old flies. With this decrease in cell number we expected a corresponding decrease in total number of epithelial nuclei, but strikingly found that number of epithelial nuclei remained constant at ~188 nuclei (Figure 2D). The nuclear morphology in aging cells was indistinguishable from those in young flies, suggesting there was no apoptosis (Figure 2A and 2B, see DAPI image). In addition, the epithelial nuclei remained diploid in 40d old flies indicating the nuclear DNA content did not change with age (Figure 2E).

**Figure 2.**
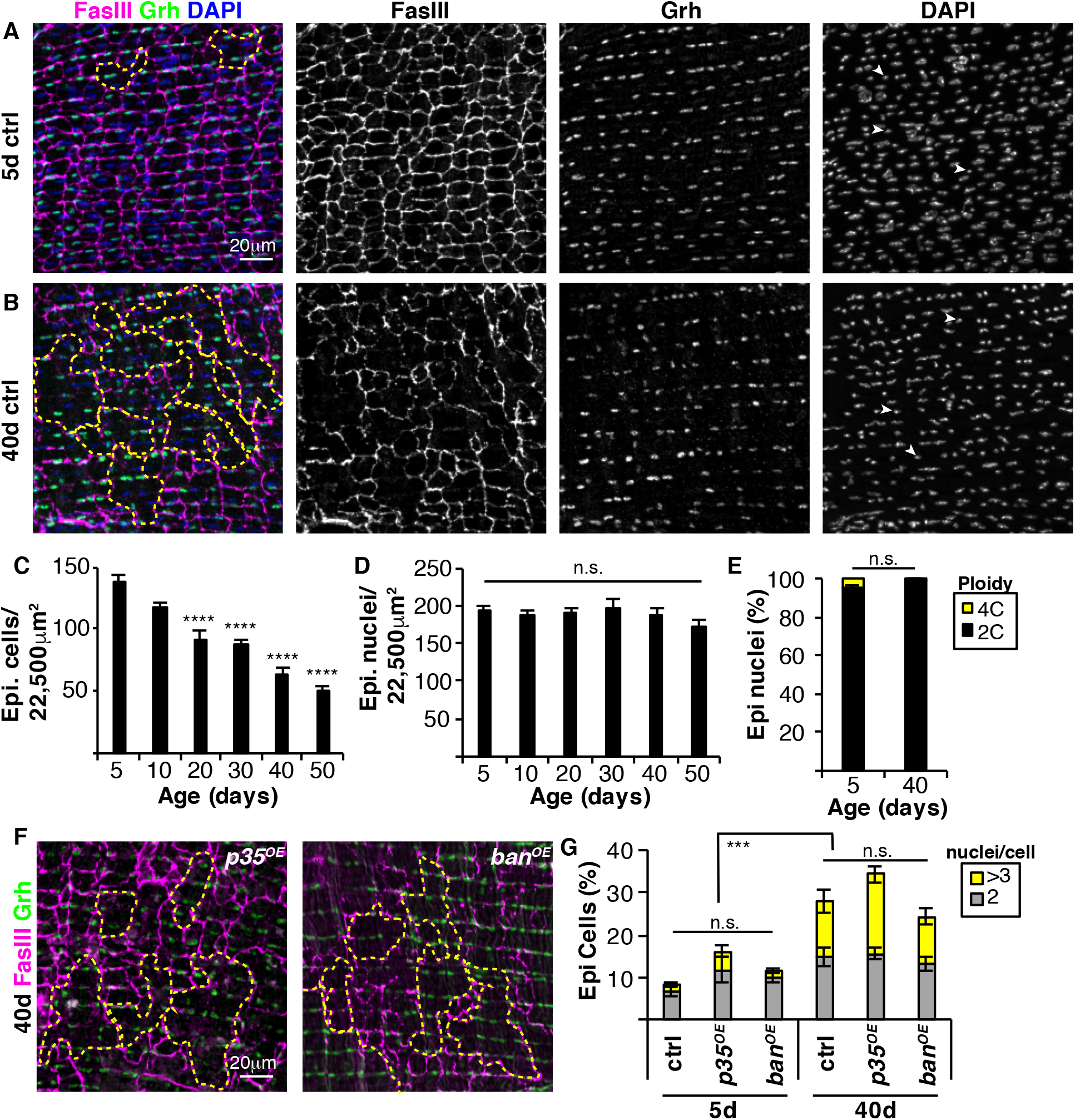
Age-induced polyploidy is not dependent on apoptosis. (A-B) Representative immunofluorescent images of 5d and 40d old female flies. Septate junctions (FasIII, magenta), epithelial nuclei (Grh, green), DAPI (blue), and multinucleated cells (outlined, yellow dashed line). (C) Epithelial cell number declines with age. (D) Number of epithelial nuclei does not change with age. (E) Quantification of epithelial nuclear ploidy (n=3 flies/ age). (F) Representative immunofluorescent images of 40d old flies with epithelium expressing anti-apoptotic genes: *p35^OE^* and *ban^OE^*. Septate junction (FasIII, magenta), epithelial nuclei (Grh, green), and multinucleated cells (outlined, yellow dashed line). (G) Percentage of bi- and multinucleated epithelial cells with age. Data represent the mean ±SE with two-way ANOVA with Šídák’s multiple comparisons test.

To further assess the role of cell death, we overexpressed cell survival genes to determine whether inhibition of apoptosis could reduce generation of polyploid cells with age. Two caspase pathway inhibitors, the gene *p35* (*p35^OE^*) and microRNA *bantam* (*ban^OE^*) were overexpressed using epi-Gal4 *Drosophila* strain (Kester and Nambu, 2011; Thompson and Cohen, 2006). However, overexpression of these caspase inhibitors did not inhibit age-induced polyploidy (Figure 2F). We still observed a significant increase in formation of multinucleated epithelial cells by 40d with no significant difference between the control flies (Figure 2G). In conclusion, cell death via apoptosis does not appear to be necessary for age-induced polyploidy in *Drosophila.*

### Polyploid cells arise by cell fusion, not endomitosis

The observed multinucleated cells in aging *Drosophila* can arise by endomitosis or cell fusion (Bailey et al., 2021; Peterson and Fox, 2021). Endomitosis occurs when cells enter the cell cycle, but fail to complete M phase. In particular, failed cytokinesis would generate a binucleated cell or multiple endomitotic cell cycles could generate a multinucleated cell. Alternatively, two or more cells can fuse together to generate bi- and multinucleated cells similar to development of myotubes that make up the *Drosophila* musculature (Lee and Chen, 2019).

To elucidate the mechanism of polyploid generation with age, we knocked down the mitotic regulators *cdc2* (*cdk1*) and *stg* (*cdc25*) using epi-Gal4 to inhibit endomitosis and examined the epithelial organization in 5d and 40d old flies. We found that binucleated and multinucleated cells were still prominent in old flies even with expression of either *cdc2^RNAi^* or *stg^RNAi^* (Figure S2A). There was a significant increase in polyploidy with age (5d vs 40d), but no significant difference in generation of polyploid cells between control and M phase gene knockdowns (Figure S2B). Likewise, the 40d old fly epithelium was still composed of ~40% multinucleated cells similar to control animals (Figure S2C), suggesting that syncytia do not arise by endomitosis.

Next, we tested whether polyploid epithelial cells arise by cell fusion. We used epi-Gal4 to express UAS-dBrainbow, a Cre recombinase-based fluorescent labeling technique to mosaically label the *Drosophila* epithelium. The Cre based recombination of the dBrainbow cassette results in expression of EGFP-HSV, EBFP2-HA, or mKO2-myc, (Hampel et al., 2011). We omitted mKO2-myc fluorescence, so we could co-stain for FasIII and validate the syncytia boundaries. Bicolored cells co-expressing both EGFP-HSV and EBFP2-HA would indicate of cell fusion (Figure 3A). In young 5d old fly tissue expressing both markers, we observed distinct patches of EGFP-HSV and EBFP2-HA with only an occasional bilabeled cell (Figure 3B and 3C). Whereas in 40d old flies, there was extensive overlap of EGFP-HSV and EBFP2-HA fluors indicative of cell fusion. Young flies had on average 1.5 ± 1.0 bicolored cells, whereas old flies had 16.9 ± 3.0 bicolored cells per epithelial area (Figure 3C).

**Figure 3.**
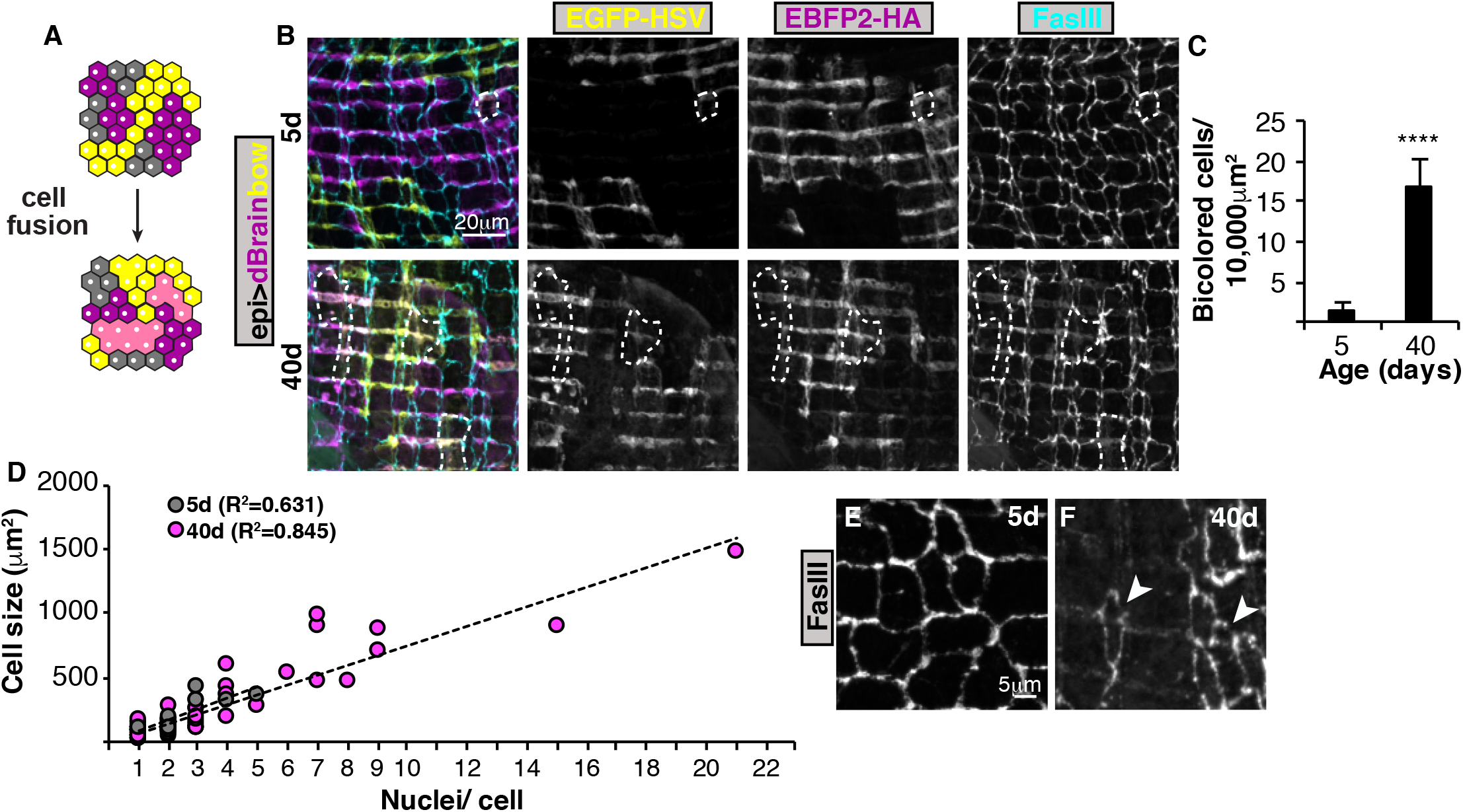
Age-induced polyploidy arises by cell fusion. (A) Illustration of mosaic labeled cells, where cell fusion leads to bicolored cells. (B) Representative images of 5d and 40d dBrainbow flies. Cells expressing both EGFP-HSV and EBFP2-HA are outlined with white dashed lines. (C) Bicolored cells arise with age. (D) Bicolored cell size and nuclear number per cell based on epithelial FasIII borders. (E and F) Representative immunofluorescent images of 5d and 40d epithelium. Some mononucleated cells in the old epithelium have signs of FasIII border deterioration (arrowheads). Also, see Figures S2.

We then examined whether bilabeled regions matched the FasIII boundaries containing more than one nucleus. At 5d, only 10 bilabeled cells were identified in 3 out of the 10 flies assayed and all but one cell had 2 or more nuclei as expected (Figure 3D). In addition, we found that cell size modestly scaled with the number of nuclei per cell at 5d (R^2^=0.63). At 40d, a total of 84 bilabeled cells were identified by their FasIII boundaries in the 10 flies assayed. Sixty percent (50 out 84 cells) had more than 2 nuclei and cell size scaled proportional with the number of nuclei per cell (Figure 3D, R^2^=0.85). Surprisingly, we found that 40% of bilabeled epithelial cells were mononucleated. We frequently observed deterioration of sepatate junctional signal in old flies, suggesting that the boundaries are in process of breaking down with age (Figure 3E and 3F). The cell junctions were scored as positive if at least 50% of FasIII signal was still present. The mononucleated cells often had breaches in FasIII border surrounding larger syncytia, suggesting these cells were in the process of fusing (Figure 3F, arrowheads). The dBrainbow bilabeling of these mononucleated cells in the old flies suggests we are underestimating the extent of cell fusion and that it is a primary mechanism for multinucleation in aging epithelium.

### Epithelial multinucleation is dependent on alpha catenin in both *Drosophila* and mice

Epithelial multinucleation has also been observed in the human and mouse RPE with normal aging as well as macular diseases (Chen et al., 2016; Saksens et al., 2016; Zhang et al., 2019). Butterfly-shaped pigment pattern dystrophy, a rare human disease (OMIM #608970), is characterized by deposits at the level of the RPE in the macular region that resemble the wings of a butterfly. Heterozygous missense mutations in alpha catenin (*Ctnna1*) are associated with this disease and multinucleation in RPE of *Ctnna1^Tvrm5^* mice (Saksens et al., 2016). In 4-month old C57BL/6J mice, the wildtype RPE is made up of centrally located, predominantly binucleated cells with only 4% multinucleated RPE cells (Figure 4A and 4C). Whereas in case of *Ctnna1^Tvrm5^* mouse mutant, the RPE is made of 21% multinucleated cells, which covers 51% of the central epithelium (Figure 4B-4D). To determine whether this is analogous to our *Drosophila* model, we knocked down alpha catenin (*αcat^RNAi^*) in fruit fly epithelium to determine if multinucleation could be induced. αCat is expressed at a low, but detectable level in adult fly epithelium and can be efficiently knocked down within 7 days of RNAi expression (Figure S3). Indeed, we found a ~4-fold increase in multinucleated epithelial cells with genetic knockdown of αcat in fly epithelium suggesting that a similar mechanism leads to multinucleation in flies and mice (Figure 4E, 4F and 4I).

**Figure 4.**
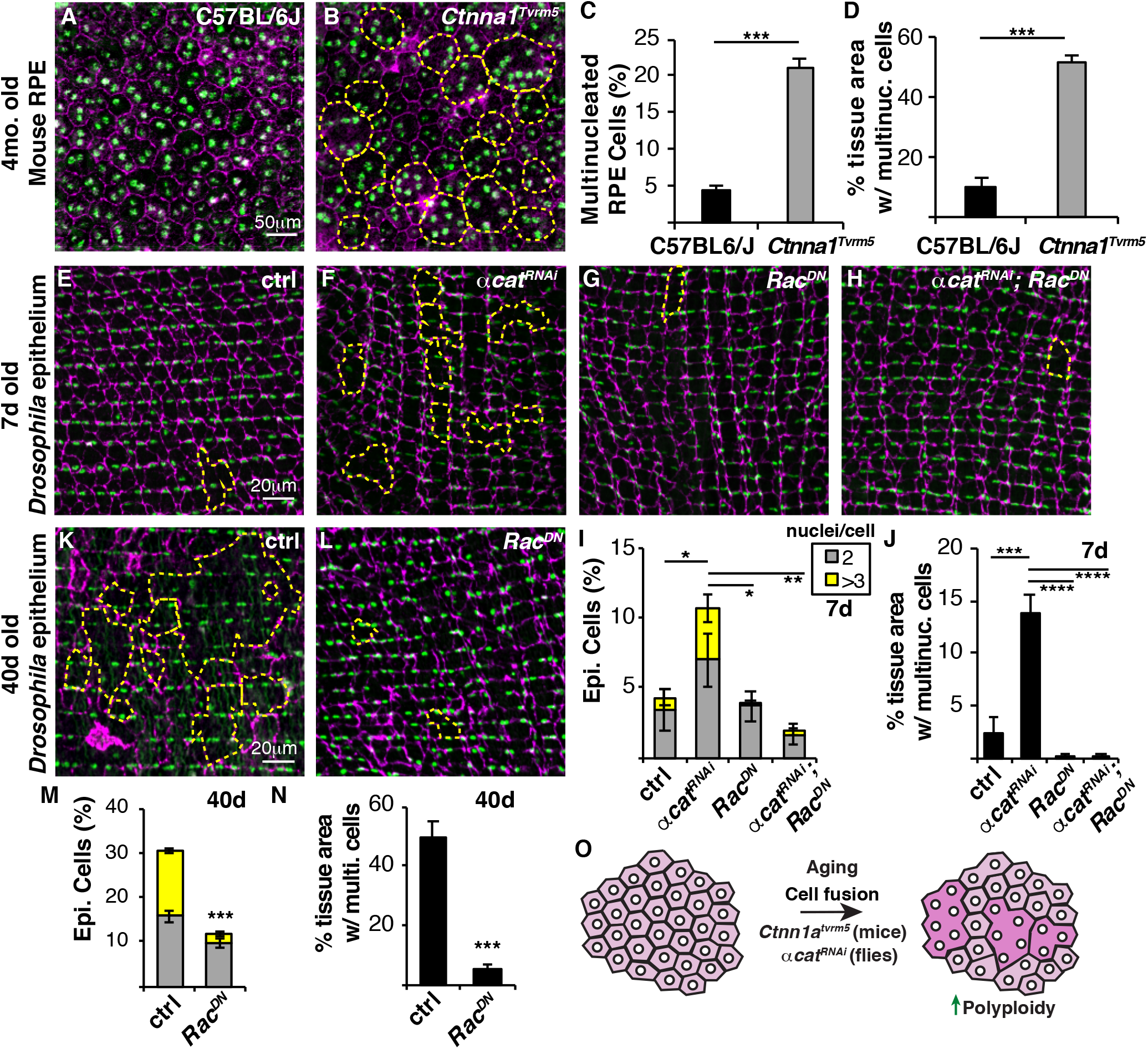
Disruption of alpha catenin causes multinucleation and inhibition of cell fusion maintains epithelial organization with age. Multinucleation is a hallmark of the RPE in the mouse model of Butterfly-shaped pigment pattern dystrophy. (A and B) Representative immunofluorescent images of RPE in wild-type (C57BL/6J) and *Ctnna^Tvrm5^* mutant mice. Cell borders (Phalloidin, magenta), RPE nuclei (DAPI, green), and multinucleated cells (outlined, yellow dashed line). (C) Percentage of multinucleated RPE cells in 4-month old mice. (D) RPE tissue area composed of multinucleated cells. (E-H) Representative immunofluorescent images of 7d epithelium from control, *αcat^RNAi^*, *Rac^DN^*, and *αcat^RNAi^*, *Rac^DN^* Drosophila strains. Septate junctions (FasIII, magenta), epithelial nuclei (Grh, green), and multinucleated cells (outlined, yellow dashed line). (I) Percentage of bi- and multinucleated epithelial cells in 7d old flies (n=5). (J)Epithelial tissue area composed of multinucleated epithelial cells in 7d old flies (n=5). (K and L) Representative immunofluorescent images of 40d old control and *Rac^DN^* flies. (M) Percentage of bi- and multinucleated epithelial cells at 40d. (N) Epithelial tissue area composed of multinucleated cells at 40d. (O) Model showing that polyploid cells arise with age due to cell fusion or inhibition of alpha catenin, a gene disrupted in patterned dystrophy. Blocking of cell fusion is sufficient to inhibit age-induced polyploidy and maintain a youthful epithelial organization even in older animals. Also, see Figures S3.

Cytoskeletal remodeling is known to play a key role in cell fusion, particularly in muscle development, which can be inhibited by expression of dominant negative Rac GTPase, *Rac^DN^* (Fernandes et al., 2005; Losick et al., 2013). We found that co-expressing *αcat^RNAi^* and *Rac^DN^* in fly epithelium significantly reduced multinucleation and the tissue area composed of multinucleated cells at 7d, indicating that genetic knockdown of *αcat* results in ectopic cell fusion (Figure 4E-4J).

### Inhibition of cell fusion prevents age-induced polyploidy and maintains epithelial organization in old animals

To maintain tissue function, one strategy is to preserve the youthful tissue organization with age. We noticed that overexpression of *Rac^DN^* appeared to reduce the epithelial tissue area composed of multinucleated cells even in young 7d old flies (Figure 4G and Figure 4J). This finding led us to ask whether blocking cell fusion would preserve epithelial organization with age. Indeed, the continuous overexpression of *Rac^DN^* in the fly epithelium was sufficient to strongly reduce generation of polyploid cells, both binucleated and multinucleated cells were significantly reduced in 40d old flies (Figure 4K-4N). Strikingly, the 40d old epithelium expressing *Rac^DN^* resembled the young 7d old control epithelium with preservation of mononucleated epithelial cells. Only 5% of the epithelium was composed of multinucleated cells in 40d *Rac^DN^* flies compared to the 50% in control flies (Figure 4N). We also measured epithelial nuclear ploidy to see if there was a compensatory increase in nuclear DNA ploidy, but found that mononucleated cells remained diploid (2C) in *Rac^DN^* epithelial cells (data not shown). In conclusion, we have characterized a new *Drosophila* model to study the mechanisms of age-induced polyploidy revealing a genetic strategy to maintain the youthful epithelial organization through inhibition of the generation of polyploid cells with age (Figure 4O).

## Discussion

Polyploidy has been found to increase with normal aging in a variety of animal tissues, including the brain, eye, heart, and liver (Chen et al., 2016; Ikebe et al., 1986; Matsumoto et al., 2021; Nandakumar et al., 2020; Senyo et al., 2013). Here we find that polyploid cells also arise in fruit fly epithelium with age. An advantage of using the fruit fly as a model is age-induced polyploidy arises by 20d, instead of months to years in mice and humans. What is more unique is the fly epithelial cells become multinucleated with age. This resembles the wound-induced polyploidization response we previously described, but the age-induced polyploid cells arise by cell fusion not the endocycle (Losick et al., 2013 and Losick et al., 2016).

In normal aging and age-associated diseases, multinucleated cells are a common feature of human and murine eye tissues made up of post-mitotic cells. The cornea endothelium is terminally differentiated and becomes multinucleated with age and the age-associated disease Fuchs endothelial corneal dystrophy (Ikebe et al., 1986; Losick et al., 2016). Likewise, the RPE is made up of terminally differentiated cells that become multinucleated with normal aging and macular diseases (Chen et al., 2016; Saksens et al., 2016; Zhang et al., 2019). In C57BL/6J mouse model, aging caused a decrease in RPE cell number and increase in cell size characterized by multinucleated cells as early as 6 months (Chen et al., 2016). However, the total number of RPE nuclei did not change with age leading to the conclusion that the multinucleated RPE cells arose by failed cytokinesis. We also observed that total epithelial nuclear number did not change with age in the fly epithelium, but it was not an indication of mechanism of cellular multinucleation. To date no study in the eye has directly assayed for cell fusion or endomitosis (failed cytokinesis) *in vivo*, so the mechanism of epithelial multinucleation remains unknown.

We show here that multinucleation in *Drosophila* epithelium is dependent on cell fusion using cell-cell junctions and mosaic dBrainbow labeling. The mosaic labeling revealed that cell fusion events may be more widespread within this tissue as there were frequent breaches in septate junctional staining that may account for the bicolored mononucleated cells observed. Alternatively, cytoplasmic sharing through the opening of gap junctions could also account for the bicolored, mononucleated cells we observed with age (Peterson et al., 2020). However, cytoplasmic sharing occurs independently of actin remodeling and we found that epithelial multinucleation was dependent on Rac GTPase.

Remodeling of the actin cytoskeleton is critical for many cell-cell fusion events from fertilization to muscle development (Brukman et al., 2019). In *Drosophila* muscle, actin polymerization in the attacking myoblast cell propels membrane protrusions into the founder muscle cell facilitating the close contact required to initiate a fusogenic synapse (Kim and Chen, 2019). Cell fusion also requires mechanical forces, including pushing and resisting cellular forces. In particular, two mechanotransduction proteins, Myosin II and Spectrins, accumulate at fusogenic synapse to mediate myoblast fusion (Duan et al., 2018; Kim et al., 2015). We still do not know why epithelial cells fuse with age and age-associated disease, but we can hypothesize that is might be related to same mechanical forces that regulate muscle cell fusion. In epithelium, α-Catenin is required to link cadherins to the actin cytoskeleton. In mice, the *Ctnna1^Tvrm5^* mutant is associated with a missense Leu436Pro substitution that maps to the proposed force-sensing module in α-catenin 1, which may be sensitive to tension (Leckband and de Rooij, 2014; Saksens et al., 2016). However, a direct link between α-Catenin, tension, and cell fusion has yet to be determined.

Still the central unanswered question is whether epithelial multinucleation is an adaptation or precursor to tissue degeneration. The key is to inhibit polyploidy and determine its long-term impact. Here we find a genetic strategy to prevent epithelial multinucleation with age by expressing *Rac^DN^* in *Drosophila* epithelium. As a result, the youthful, mononucleated epithelial organization can be maintained. Future studies will then aim to elucidate if epithelial physiology is improved by inhibition of age-induced polyploidy.

## Materials and Methods

### Fly husbandry and strains

*Drosophila melanogaster* strains used in this study were reared on standard corn syrup-soy food (Archon Scientific) at 25°C, 60% humidity, and 12h light/ dark cycle. The following *Drosophila* strains were obtained from Bloomington (b) or created in lab using the strains indicated: GMR51F10-Gal4, referred to as epi-Gal4 (b38793), *w^1118^* (b3605), CantonS (b64349), UAS-*cdc2^RNAi^* (b28368), UAS-*stg^RNAi^* (b29556), UAS-*Rac^DN^* (b6292), UAS*-αcat^RNAi^* (b38985), UAS-dBrainbow (b34513), Crey/FM7;; 51F10-Gal4/TM3 (derived from epi-Gal4 (b38793), Crey (b851), and Sn28/FM7;; TM2/TM6b balancer (obtained from Dr. Tina Tootle, University of Iowa), UAS-*αcat^OE^* (b58787), UAS-*p35^OE^* (b5073), and UAS-*ban^OE^* (b60672). All flies used in this study were female and the control, unless otherwise noted, was the heterologous epi-Gal4/*w^1118^* strain. Epi-Gal4/UAS system was used to drive gene expression or RNAi knockdown unless otherwise noted.

### Mouse husbandry and strains

Mouse strains used in this study were C57BL/6J-*Ctnna1^Tvrm5^*/PjnMmjax and C57BL/6J (The Jackson Laboratory, stock #43572 and #000664, respectively) were bred and maintained under standard conditions of 12:12 light-dark cycle in the Research Animal Facility at The Jackson Laboratory. Mice were provided with NIH31 (6% fat chow) diet and HCl acidified (pH 2.8-3.2) water ad libitum and maintained in pressurized individual ventilation cages, which were regularly monitored to maintain a pathogen-free environment. The C57BL/6J-*Ctnna1^Tvrm5^*/PjnMmjax mice were maintained on C57BL/6J background and confirmed for the absence of *Crb1^rd8^* mutation.

### *Drosophila* aging, dissection, and immunostaining

Newly eclosed females, aged as denoted in figure legend, were dissected as previously described (Bailey et al., 2020). Briefly, abdomens were fixed in 4% paraformaldehyde, permeabilized in 1x PBS with 0.3% Triton-X-100 and 0.3% BSA, then stained overnight at 4°C using primary antibodies. In this study, mouse anti-FasIII (DSHB, 7G10, AB_528238, 1:50), rabbit anti-Grh (AB_2568305, 1:300) (Losick et al., 2016), rabbit anti-GFP (Thermofisher, A-11122, AB_221569, 1:2000), rat anti-HA (Roche, 11867423001, AB_390918, 1:100), and mouse alpha-catenin (DSHB, D-Cat, AB_532377) primary antibodies were used. Secondary antibodies from Thermofisher included donkey anti-rabbit 488 (A21206, AB_2535792), goat anti-mouse 568 (A11031, AB_144696), and goat anti-rat 633 (A21094, AB_2535749) used at 1:1000 dilution. All tissues were stained with DAPI at 10μg/mL and mounted in Vectashield (Vector Laboratories, H1000-10) on a glass coverslip and slide, with the inner tissue facing out.

### *Drosophila* imaging and analysis

Abdomens were imaged using a Zeiss ApoTome and a 40x dry objective. Z-stack images were taken at 0.50-0.55μm per slice. Using the NIH FIJI software (Schindelin et al., 2012), images flattened using SUM slices tool for DAPI images and MAX projection for other channels.

**Cell area** was measured using FasIII (septate junctions) to determine cell boundaries. An area of 100×100μm or 150×150μm at least 25μm away from the dissected edge was chosen. The image was converted to a binary black/white image using the threshold tool, and any missing segments of cell boundary as determined by the analyzer while comparing the thresholded image and the original FasIII image were drawn in using the pencil tool set at 3μm. Any portion of the image that did not have a FasIII signal at least twice the brightness of the background was not considered a border. Areas of staining more than 3μm apart were not considered a continuous border. A region of interest (ROI) map was generated using the analyze particles tool, which was used to calculate the cell area. Any cells that extended off the edges of the area being analyzed were excluded. For each cell, the number of nuclei per cell was found by merging the FasIII and Grh (epithelial nuclear marker) channels and counting the number of nuclei present in each cell.

**Area covered by multinucleated cells** was calculated by summing the areas of each cell with 3 or more nuclei and dividing by the total area of all cells measured in a given image, then multiplying by 100.

**dBrainbow** mosaic labeling was used to quantify the number of fused cells per 100×100μm epithelial area. Bicolored regions were identified by epithelial regions co-expressing EGFP-HSV-Alexa 488 and EBFP2-HA-Alexa 633 fluorescence. Bicolor cell areas and nuclear number were further quantified by measuring the FasIII borders and DAPI stained nuclei.

**Fluorescent intensity** was measured for α-Cat protein expression. To do so, 50 regions of interest were drawn around Grh^+^ epithelial area and transferred to α-Cat channel. The integrated density of α-Cat calculated to determine the average fluorescence intensity for each *Drosophila* abdominal epithelium.

**Epithelial ploidy** was calculated according to (Bailey et al., 2020). Samples from 5d and 40d old flies were dissected and stained as described above, then imaged with the same exposure time. In FIJI, ROIs were drawn around each nucleus in the Grh (epithelial nuclear marker) channel. This ROI map was superimposed on the sum of slices of the Z-stack for the DAPI channel. Nuclei that overlapped with fat body or other tissue were excluded, then the area and integrated density was measured for each nucleus. The intensity was calculated by subtracting the background, then comparing the intensity of nuclei from old flies to those of the young flies whose nuclei were previously shown to be diploid (Losick et al., 2013).

***Drosophila* Lifespan** was determined by placing 10 newly eclosed females and 5 males each in 10 food vials total. The numbers of live female flies were counted each day. Flies were flipped to new food every 2-3 days, and maintained with male flies for duration of the lifespan assay.

### Mouse RPE flat mounts and immunostaining

For RPE flatmounts, 4-month-old C57BL/6J-*Ctnna1^Tvrm5^*/PjnMmjax and age-matched C57BL/6J mice were asphyxiated by carbon dioxide inhalation. The enucleated eyes were marked with an orange dot dorsally to orient the eyes and placed in ice-cold 4% paraformaldehyde (PFA) solution. The extra-ocular tissue and the anterior segment of the enucleated eyes were removed. The eye cups, which were fixed overnight in 4% PFA at 4 °C, were washed in 1X TBS and the neural retina was separated from the RPE-choroid-sclera. Six radial cuts were made toward the optic nerve to flatten the posterior eyecup. RPE cell borders were stained with Phalloidin (F-actin) and nuclei were stained by incubating with 4′,6-diamidino-2-phenylindole (DAPI) for 2 days at 4 °C with agitation. Both Phalloidin and DAPI were prepared in 0.3% Triton in 1X TBS. The RPE-choroid-sclera tissue was then moved through a glycerol gradient of 10%, followed by 20% and 50%, and allowed to equilibrate in each gradient for about 8 hours at 4 °C with constant agitation. The whole RPE-choroid-sclera tissue was then flattened, with RPE side up, onto slides, mounted in Vectashield (Vector Laboratories) overlaid with a cover slip, and examined under a fluorescent Zeiss Axio Observer.Z1 Microscope (Carl Zeiss AG). Identical imaging parameters were applied to both *Ctnna1^Tvrm5^* and C57BL/6J control RPE flat mounts. For cell and nuclear count, identical regions were selected in oriented eyes from both strains, and the cells and nuclei within those cells were counted by selecting a uniform area of 350μm^2^ for each image, using the same protocol as described above for the *Drosophila* cell and nuclear count, in FIJI software.

### Replicates and Statistical Analysis

All *Drosophila* experiments analyzed contained a minimum of 10-15 biological replicates (fruit flies) obtained from 2 or more experiments. Mouse RPE analysis used 3 biological replicates (mice). Data represents the mean ± standard error and statistical analysis was performed with GraphPad Prism 9. T-test with Welch’s correction was used for pairwise analysis and two-way ANOVA with Tukey’s multiple comparisons test unless otherwise noted in the Figure Legend. P-values are denoted as ns (p>0.05), * (p<0.05), ** (p<0.01), *** (p<0.001), and ****(p<0.0001).

## Acknowledgements

We would like to thank the members of the Losick lab (Dr. Erin Bailey, Levi Duhaime, Meaghan Grogan, and Minqi Shen), for critical review of this manuscript and the fly community, particularly the Bloomington Drosophila Stock Center (NIH P40OD018537), the Developmental Studies Hybridoma Bank (created by the NICHD and maintained at The University of Iowa, Department of Biology, Iowa City, IA 52242, USA) and the TRiP Center at Harvard Medical School (NIH/NIGMS R01-GM084947) for providing transgenic stocks or additional reagents used in this study. Images were acquired using equipment of the Light Microscopy Facility at the MDI Biological Laboratory, which is supported by the Maine INBRE grant (GM103423) from the National Institute of General Medical Sciences at the National Institutes of Health as well as Bret Judson and the Boston College Imaging Core for infrastructure and support. Research reported in this publication was supported by MDI Biological Laboratory, Procter Award to V.P.L., Boston College, and the National Institute of General Medical Sciences of the National Institutes of Health under Award Number R35GM124691 to V.P.L. and the National Eye Institute under EY027860 and EY011996 to P.M.N.

## Author Contributions

Conceptualization, A.S.D., N.G., and V.P.L.; Methodology, A.S.D., N.G., P.M.N., and V.P.L.; Validation, A.S.D. and N.G.; Formal Analysis, A.S.D., N.G., and V.P.L.; Investigation, A.S.D. and N.G.; Resources, A.S.D., N.G., P.M.N., and V.P.L.; Writing – Original Draft, A.S.D., N.G., and V.P.L.; Writing – Review & Editing, A.S.D., N.G., P.M.N., and V.P.L.; Visualization, A.S.D., N.G., and V.P.L.; Project Administration, V.P.L.; Supervision, P.M.N., and V.P.L.; Funding Acquisition, P.M.N. and V.P.L.

## Declaration of Interests

The authors declare no competing interests.

## Notes

### Competing Interest Statement

The authors have declared no competing interest.

