## Supplementa Materials and Figures for "Blocking cell fusion inhibits age-induced polyploidy and maintains epithelial organization in *Drosophila*"

**Supplemental Figure Legends**

**Figure S1. Multinucleated epithelial cells arise with age in *Drosophila melanogaster* strains.** (A-D) Representative immunofluorescent images of 5 and 40d old $w^{1118}$ and CantonS flies. Septate junction (FasIII, magenta), epithelial nuclei (Grh, green), and multinucleated cells (outlined, yellow dashed line). (E) Percentage of bi- and multinucleated epithelial cells. (F) Epithelial tissue area composed of multinucleated cells and analyzed two-way ANOVA with Šídák's multiple comparisons test.

**Figure S2. Age-induced polyploidy is not dependent on endomitosis.** (A) Representative immunofluorescent images of 40d old control, $cdc2^{RNAi}$ and $stg^{RNAi}$ flies. Septate junction (FasIII, magenta), epithelial nuclei (Grh, green), and multinucleated cells (outlined, yellow dashed line). (B) Percentage of bi- and multinucleated epithelial cells (n=5 flies/age). (C) Epithelial tissue area composed of multinucleated cells (n=5 flies/age) and analyzed with two-way ANOVA with Šídák's multiple comparisons test.

**Figure S3. Validation of alpha catenin RNAi in *Drosophila*.** (A-C) Representative immunofluorescent images of control, $\alpha cat^{RNAi}$, and $\alpha cat^{OE}$ flies. αCat (magenta) and epithelial nuclei (Grh, green). Arrowheads denote epithelial cells expressing αCat. (D) Quantification of αCat fluorescent intensity at 7d (n=9 flies per condition). Data represent the mean ± SE with unpaired T-test.

#### Figure S1

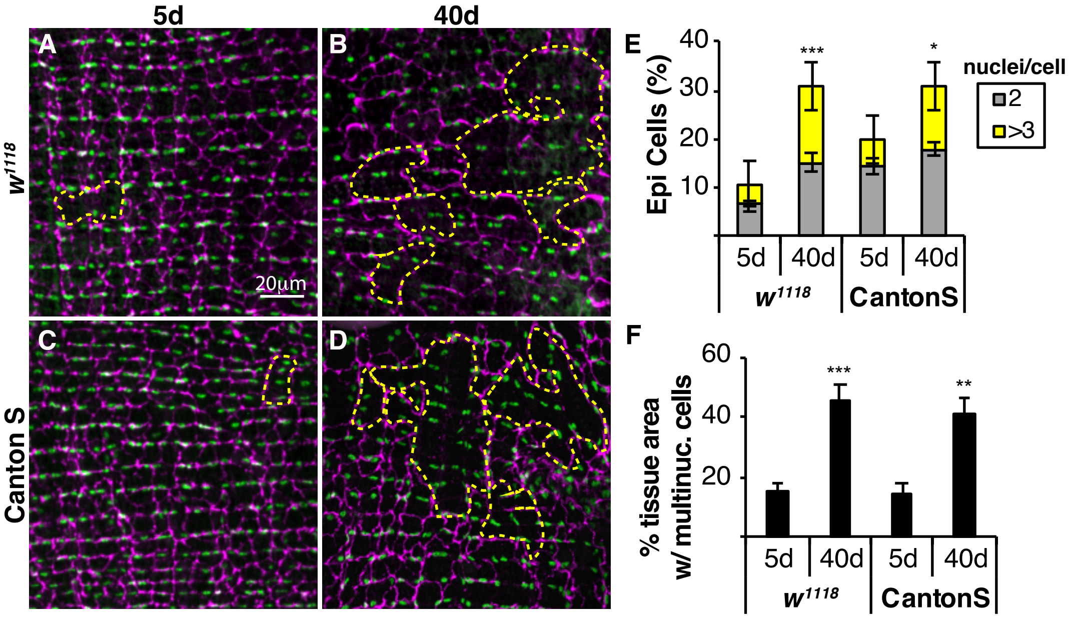

5d
40d
A
B
C
D
w1118
Canton S
20μm
E
Epi Cells (%)
40
30
20
10
***
*
nuclei/cell
2
>3
5d
40d
5d
40d
w1118
CantonS
F
% tissue area
w/ multinuc. cells
60
40
20
***
**
5d
40d
5d
40d
w1118
CantonS

### Figure S2

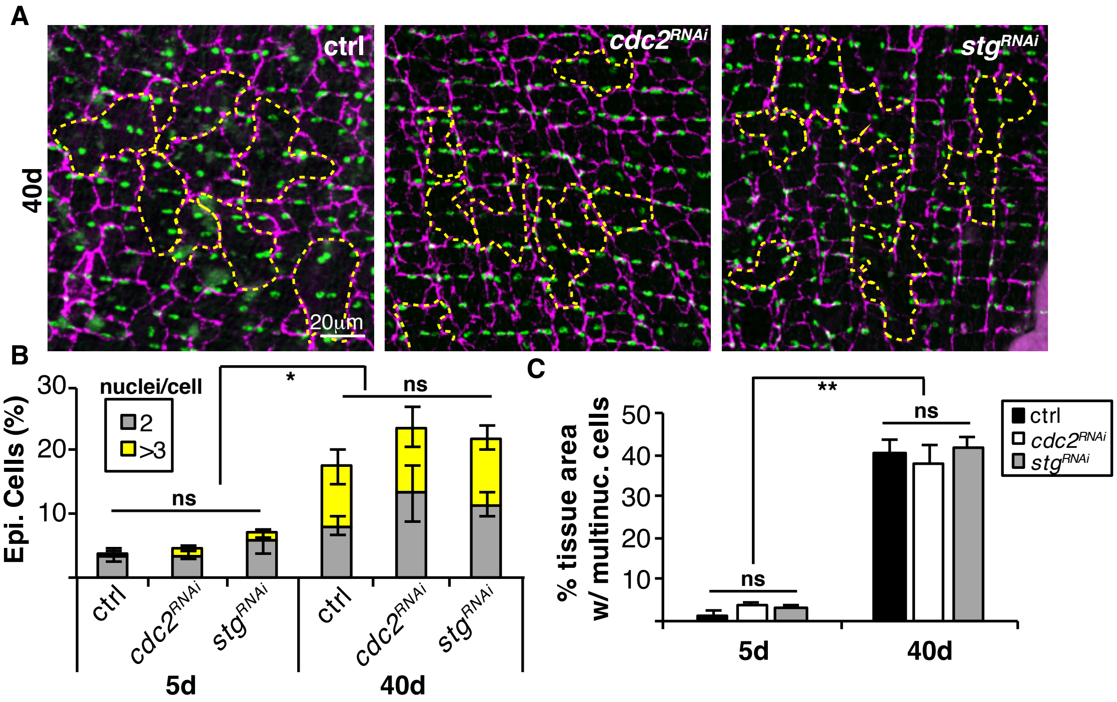

A
40d
ctrl
cdc2RNAi
stgRNAi
20μm
B
nuclei/cell
2
>3
Epi. Cells (%)
30
20
10
ns
*
ns
ctrl
cdc2RNAi
stgRNAi
ctrl
cdc2RNAi
stgRNAi
5d
40d
C
% tissue area w/ multinuc. cells
50
40
30
20
10
ns
**
ns
ctrl
cdc2RNAi
stgRNAi
5d
40d

### Figure S3

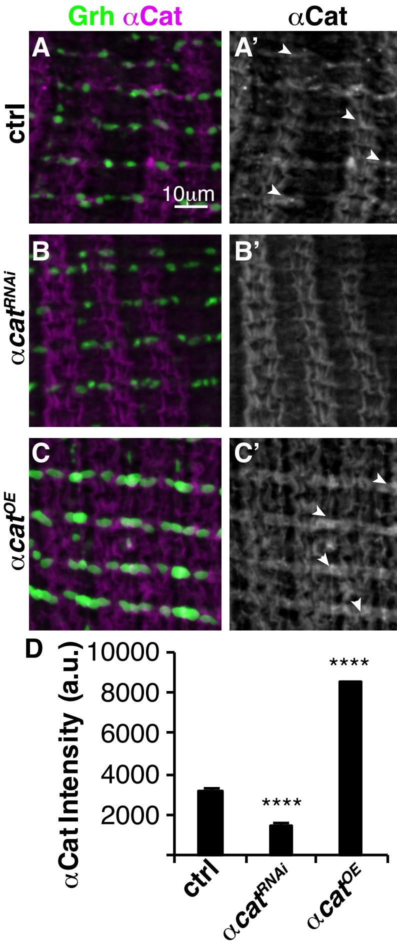
Grh αCat
αCat
A
A'
ctrl
10μm
B
B'
αcat^RNAi
C
C'
αcat^OE
D
αCat Intensity (a.u.)
10000
8000
6000
4000
2000
****
****
ctrl
αcat^RNAi
αcat^OE
